# Cryo-EM images of phase separated lipid bilayer vesicles analyzed with a machine learning approach

**DOI:** 10.1101/2023.12.31.573132

**Authors:** Karan D. Sharma, Milka Doktorova, M. Neal Waxham, Frederick A. Heberle

## Abstract

Lateral lipid heterogeneity (i.e., raft formation) in biomembranes plays a functional role in living cells. Three-component mixtures of low- and high-melting lipids plus cholesterol offer a simplified experimental model for raft domains in which a liquid-disordered (Ld) phase coexists with a liquid-ordered (Lo) phase. Using such models, we recently showed that cryogenic electron microscopy (cryo-EM) can detect phase separation in lipid vesicles based on differences in bilayer thickness. However, the considerable noise within cryo-EM data poses a significant challenge for accurately determining the membrane phase state at high spatial resolution. To this end, we have developed an image processing pipeline that utilizes machine learning (ML) to predict the bilayer phase in projection images of lipid vesicles. Importantly, the ML method exploits differences in both the thickness and molecular density of Lo compared to Ld, which leads to improved phase identification. To assess accuracy, we used artificial images of phase-separated lipid vesicles generated from all-atom molecular dynamics simulations of Lo and Ld phases. Synthetic ground truth datasets mimicking a series of compositions along a tieline of Ld+Lo coexistence were created and then analyzed with various ML models. For all tieline compositions, we find that the ML approach can correctly identify the bilayer phase at 5 nm lateral resolution with > 90% accuracy, thus providing a means to isolate the intensity profiles of coexisting Ld and Lo phases, as well as accurately determine domain size distributions, number of domains, and phase area fractions. The method described here provides a framework for characterizing nanoscopic lateral heterogeneities in membranes and paves the way for a more detailed understanding of raft properties in biological contexts.

**Significance:** Lipid rafts are important for cell function, but in most cases cannot be detected with conventional optical microscopy because of their extremely small size. Cryogenic electron microscopy (cryo-EM), because of its much greater spatial resolution, is capable of imaging domains as small as 5-10 nm. In this report, we show how machine learning techniques can be used to automatically and accurately identify raft-like domains in simulated cryo-EM images, a powerful approach that could ultimately lead to a better understanding of raft properties.

## 1. Introduction

The membranes that surround and partition living cells can play both structural and functional roles. The latter is exemplified by the lipid raft hypothesis, which proposes the existence of plasma membrane (PM) domains enriched in sphingolipids and cholesterol that are thought to direct the spatial organization of membrane proteins and thus mediate processes such as cell signaling (1). Multiple lines of evidence have converged on a picture of raft domains as small (< 20 nm) and highly dynamic in resting conditions, but with the ability to coalesce into larger structures in response to certain stimuli (2). The structural and elastic properties of the raft and non-raft environments can differ substantially, with the latter being thinner, easier to bend, less viscous, and more compressible owing to a greater proportion of unsaturated lipids (3–5). These local membrane properties influence the free energy of protein conformations (6) and can result in distinct phase preferences for different proteins (7). In the raft model of PM organization, the size, connectivity, and composition of both raft and non-raft domains are all variables that can affect the local concentrations of membrane proteins and thus, the frequency of encounters between interaction partners.

One of the great challenges in studying lipid rafts is their small size, which precludes direct observation by conventional optical microscopy. Instead, researchers have relied heavily on biochemical and spectroscopic data to infer the presence or absence of multiple membrane environments in cell membranes. While the lack of visual evidence initially provided fuel for controversy, those concerns have been largely put to rest by the sheer quantity of indirect data that consistently points toward heterogeneous membranes as a rule rather than an exception (8). Yet even as the “seeing is believing” debate recedes into the background, a fact often lost in the discussion is that the utility of image data extends far beyond visual confirmation of rafts. To take one example, images can reveal spatial domain patterns that might help distinguish between competing theories of raft formation (9). Image data also open a window to heterogeneity in domain structures that are often obscured in ensemble-averaged spectroscopic or scattering measurements, thus providing a more complete picture of rafts and their role in specific biological processes.

The great wealth of information contained in sub-optical images is exemplified by the emergence of single-particle cryo-electron microscopy (cryo-EM) as a pre-eminent tool in structural biology. A crucial breakthrough that has accelerated its rise is the use of machine learning (ML) at multiple stages of the experimental workflow (10) including image denoising (11), particle picking (12), image segmentation (13), and 3D reconstruction (14,15). Far from being limited to water-soluble proteins, cryo-EM is increasingly being used to determine structures of membrane proteins in native or near-native environments (16–20). As the focus of these studies has been fixed squarely on the protein, their membrane hosts have received less attention, yet bilayers also appear in images at a resolution high enough to observe density variation in the separate leaflets. This raises the possibility that, when visualized with cryo-EM, relatively thicker raft domains might be distinguishable from the thinner “sea” of mostly unsaturated lipids that surrounds them.

To this end, we recently developed a method for obtaining local intensity profiles along the projected circumference of lipid vesicles in cryo-EM images (21), which we then used to measure the local bilayer thickness at 5 nm lateral resolution in liposomes and vesicles derived from cell plasma membranes (21) as well as HIV pseudoviral membranes (22). When thicknesses from a large population of vesicles were plotted as a histogram, the appearance of the distribution (i.e., uni- or multi-modal) was consistent with the number of coexisting phases determined in independent experiments. Concurrently with our study, the Keller group performed similar analyses and reached similar conclusions using tomographic reconstructions of cryo-preserved liposomes (23). These outcomes were expected, since it is well established that ordered phases are 5-10 Å thicker than disordered phases (3,24,25). However, while those studies demonstrated critical proof-of-principle for using cryo-EM to investigate lipid phase separation, its unique capabilities as an imaging technique have not yet been fully realized. Most importantly, the development of a robust, automated method for determining the membrane phase state with high spatial resolution would enable additional valuable characterization including domain size distributions and an assessment of heterogeneity within the sample (26).

Although the local phase state of the membrane can in principle be determined from spatially resolved bilayer thickness measurements, in practice this is hampered by the substantial noise inherent to cryo-EM images. However, bilayer thickness is not the only characteristic that distinguishes ordered and disordered phases. Equally important from the perspective of electron scattering in image formation is the difference in molecular packing density, which manifests as a greater intensity contrast for ordered phases compared to disordered phases (27). Here, we show that ML models trained to recognize both thickness and intensity contrast can predict the local bilayer phase state with significantly greater fidelity than methods based on thickness measurements alone. Key to this conclusion is the use of *in silico* data that allows us to establish a ground truth for assessing the accuracy of phase determination for a variety of methods, including both unsupervised and supervised ML techniques. We also describe additional structural information that can be gleaned from segmented images such as phase area fractions, average intensity profiles of ordered and disordered phases, and measurements of domain size and number.

## 2. Methods

### 2.1 Analysis software

The analyses described in the following sections were implemented in Wolfram Mathematica 13.2 (Wolfram Research, Champaign, IL) unless stated otherwise.

### 2.2 Molecular dynamics (MD) simulations of Ld and Lo bilayers

All-atom molecular dynamics simulations of three-component mixtures containing 1,2-distearoyl-sn-glycero-3-phosphocholine (DSPC), 1,2-dioleoyl-sn-glycero-3-phosphocholine (DOPC), and cholesterol were performed with NAMD (28) using the CHARMM36 force field parameters (29,30). Two compositions were simulated, corresponding to the endpoints of a tieline in the Ld+Lo region of the room temperature phase diagram (Fig. S1). Each bilayer contained 100 lipids per leaflet, 50 waters per lipid, and no ions, and was constructed and equilibrated with the CHARMM-GUI protocols (31–33). The production runs for the two bilayers (917 ns for Ld and 1345 ns for Lo) were performed at constant temperature of 293 K and constant pressure of 1 atm using the same simulation parameters as in (34). Number and charge density profiles of each lipid and water atom in the systems were calculated from the last 480 ns of the centered bilayer trajectories with the Density Profile tool in VMD (35). The dipole potential profile for each system was calculated from the charge density profile following the approach in (36). The protocol is based on Poisson’s equation and utilizes a double integral calculation, setting the potential to be the same on the opposite sides of the simulation box, which is a good approximation for symmetric bilayers (37). Table 1 gives the compositions and some properties of the simulated Lo and Ld phase bilayers.

**Table 1.** Compositions and structural parameters of simulated Ld and Lo phases at 20°C.

| Mixture | $\chi_{DSPC}$ | $\chi_{DOPC}$ | $\chi_{CHOL}$ | $A_L$ [$\text{\AA}^2$] | $D_B$ [$\text{\AA}$] |
| --- | --- | --- | --- | --- | --- |
| Ld | 0.09 | 0.79 | 0.12 | 59.9 | 40.0 |
| Lo | 0.60 | 0.11 | 0.29 | 40.0 | 52.8 |

### 2.3 Synthetic cryo-EM images

Synthetic cryo-EM images of lipid vesicles were generated following a previously described method that we summarize here (21). In phase contrast imaging mode, the observed signal is related to variation in the electrostatic potential Φ within the lipid bilayer, which depends on lipid composition and packing density and is therefore different for Lo and Ld phases. We approximated Φ(*w*) (where *w* denotes position along the direction normal to the plane of the bilayer) as a sum of contributions from individual lipid and water atoms with an additional contribution from the membrane dipole potential Φ*_d_*, i.e.,

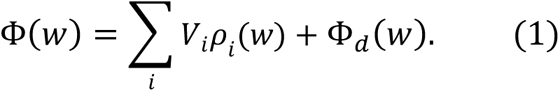

In Eq. 1, the sum is over individual lipid atoms, *ρ_i_*(*w*) is the atom’s time-averaged atomic number density profile (units of Å^-3^) calculated from the MD simulation trajectory, and *V_i_* is the spatially integrated, shielded coulomb potential for an isolated neutral atom (*V_i_* = 25, 130, 108, 97, and 267 *V*Å^3^ for H, C, N, O and P, respectively). The phase shift experienced by an electron wave passing through the bilayer is given by:

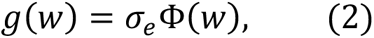

where *σ*_e_ accounts for the dependence of the electron phase on the projected potential and is equal to 0.65 *mrad V*^-1^Å^-1^for 300 keV electrons (38). Following Wang et al., we refer to *g*(*w*) as the electron ‘scattering profile’ of the flat simulated bilayer; as Φ has units of *V*, the scattering profile has units of *mrad* Å^-1^. A cryo-EM image of a liposome corresponds to a projection of the vesicle’s spherical scattering profile, *γ*(*r*, *θ*, *ϕ*), onto a plane, followed by convolution with a contrast transfer function (CTF). For a compositionally uniform liposome of radius *R*, *γ* has no angular dependence and can be approximated with the flat bilayer scattering profile *g*(*w*) using the coordinate transformation *r* = *w* + *R*, such that *γ*(*r*, *θ*, *ϕ*) = *γ*(*r*) = *g*(*r* − *R*). In this case, the projected density Γ(***r***) is easily obtained by numerical integration:

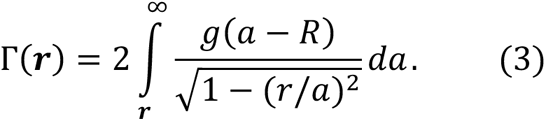

Eq. 3 is not straightforward to evaluate for a phase-separated liposome where *γ* may have a complicated angular dependence. Instead, we used a Monte Carlo technique to approximate Γ(***r***). Briefly, a three-dimensional representation of the vesicle was constructed as a set of discrete points whose spatial density was proportional to Δ*γ*(*r*, *θ*, *ϕ*) = *γ*(*r*, *θ*, *ϕ*) − *γ*_*s*_, where *γ*_*s*_ is the electron phase shift caused by vitreous ice under given imaging conditions and is equal to 3.28 *mrad* Å^-1^ here. We used a geometrical model for the vesicle in which the Ld and Lo phases each comprised a single spherical cap centered on opposite poles of the vesicle. For this domain arrangement, the position of the domain boundary is specified by the polar angle *θ_b_* = cos^−1^(1 − 2*a_Lo_*), where *a_Lo_*. is the area fraction of Lo phase. With the Lo and Ld domains centered at *θ* = 0 and *π*, respectively, the azimuthally symmetric spherical scattering profile is

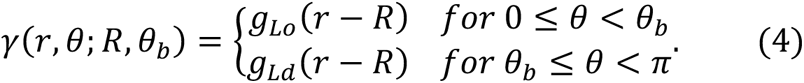

To generate a vesicle image, the vesicle radius was drawn from a Schulz distribution centered at 25 nm with a width of 5 nm, consistent with experimentally observed vesicle size distributions after extrusion through a 50 nm pore size filter. Random points were then generated within the Lo and Ld domains delineated by *θ_b_* and further subdivided radially into thin (0.2 Å-width) concentric shells, with the shell radii *r_j_* chosen to span the full thickness of the simulated flat Lo and Ld bilayers. Within the Lo and Ld shells, the point density was proportional to *γ*(*r*_*j*_, *θ*) given by Eq. 4, with the constant of proportionality chosen to generate a total of ∼10^8^ points in the vesicle. The full set of points thus generated was then randomly rotated in 3D, projected onto the *xz* plane, and binned into 2.5 Å square pixels to create the image Γ(***r***) where the intensity at each pixel was proportional to the density of projected points. At this step, the fraction of signal in each pixel originating from Lo and Ld domains was recorded and used to generate ground truth maps of the bilayer phase state in the projection image. These maps were subsequently used to assess the accuracy of machine learning analyses as described in Sections 2.6-2.7.

To mimic experimental images, Γ(***r***) must be convolved with a contrast transfer function *c*(***s***) and associated phase perturbation factor *χ*(***s***):

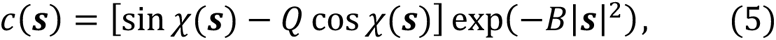

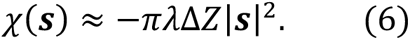

In Eqs. 5-6, ***s*** is the spatial frequency in units of Å^-1^, *B* is the amplitude decay factor in units of Å^2^, *Q* is the unitless amplitude contrast factor, *λ* is the electron wavelength, Δ*Z* is the defocus length (defined such that positive values indicated underfocus), and we have neglected spherical aberration effects in Eq. 6. The reciprocal space image is then calculated as:

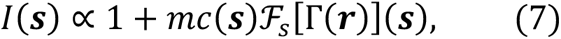

where ℱ*_S_* is the Fourier transform of the projected 2D phase shift image Γ(***r***) and *m* is an arbitrary scale factor. The corresponding real-space image *I*(***r***) is the inverse Fourier transform of *I*(***s***),

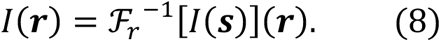

Finally, Gaussian noise was introduced into the simulated image to approximate noise in experimental images. For each simulated composition, a total of 490 vesicle images were generated.

### 2.4 Bilayer intensity profiles

Within projection images, vesicles were identified and subdivided as previously described (21) to obtain spatially resolved intensity profiles in the direction normal to the bilayer, referred to here as *I*(*w*) profiles, with *w* = 0 corresponding to the bilayer midplane. Briefly, vesicle contours (i.e., the set of points approximating the midplane of the projected bilayer as defined by a relatively bright central peak) were first generated using a neural network-based algorithm (MEMNET) developed and kindly provided by Dr. Tristan Bepler (MIT). While we retain the original MEMNET nomenclature, these algorithms (and others) are now part of the TARDIS software package (39). The MEMNET contour was resampled at arc length intervals of ∼ 5 nm, resulting in a polygonal representation of the 2D contour. For each polygon face, all pixels within a 5×20 nm rectangular region of interest centered at the face were selected, and their intensities binned at 1 Å intervals in the long dimension (i.e., normal to the face) and subsequently averaged in the short dimension to produce a local *I*(*w*) profile.

### 2.5 Bilayer thickness measurement

A key measurement obtained from *I*(*w*) is the spatially resolved bilayer thickness *D*_*TT*_, details of which are provided in previous work (21). Briefly, *D*_*TT*_ was calculated as the distance between the two minima in *I*(*w*). Two methods were used to locate the minima: (1) a ‘model-free’ method, in which a local 5-point Gaussian smoothing was first performed, and the distance between the two absolute minimum intensity values on either side of *w* = 0 was calculated; (2) a ‘model-fit’ method that fits the profile as a sum of four Gaussians and a quadratic background, with the troughs corresponding to the two absolute minimum intensity values on either side of *w* = 0. The two methods generally agree to within 1 Å, and the *D*_*TT*_ values we report are the average of the two measurements.

### 2.6 Machine learning (ML) classification of bilayer phase state

We tested various unsupervised and supervised learning methods for identifying the phase state of the bilayer, using the spatially-resolved *I*(*w*) profiles as features for discriminating Lo and Ld phases. Each profile can be considered as a 201-dimensional vector of intensities for input to the ML algorithm (where noted, the dimensionality of the input vectors was first reduced using principal component analysis). For unsupervised learning, we used *k*-means and *k*-medoids clustering with the number of clusters (*k*) set to two. For supervised learning, we used five methods as implemented in Mathematica v13.2: Decision Tree (DT), Gradient Boosted Trees (GBT), Random Forest (RF), Nearest Neighbors (NN), and Logistic Regression (LR). In every case, models were trained on *I*(*w*) profiles from pure Lo and Ld datasets and then used to predict the phase state of vesicles in which Ld and Lo phases coexist. We used 70% of the pure Ld and Lo datasets for training, with the remaining 30% reserved to assess accuracy.

### 2.7 Quantifying the accuracy of ML analysis

We assessed the accuracy of each ML model by comparing the predicted phase state to the ground truth phase state at the level of individual 5 nm segments. Because each projected bilayer segment may contain contributions from both phases, we define here the segment’s ‘ordered character’ (*OC* ∈ [0,1]) as the fraction of the segment’s intensity that originated from an Lo domain. For vesicles that contain a single domain of each phase (i.e., the geometric model used here), most segments are purely Ld (*OC* = 0) or Lo (*OC* = 1). Exceptions occur in the vicinity of the domain boundary, where intermediate *OC* values are typically observed. Importantly, all supervised ML methods yield an estimate of the probability that a given segment is Lo (i.e., a prediction of the segment’s *OC* value). We therefore used the residual (i.e., *OC* − *OC*′, where *OC* is the ground truth ordered character and *OC*′ is the prediction of the ML model) as one metric for accuracy.

Unsupervised ML methods result in a binary classification of the segment phase state (i.e., a given segment is either Ld or Lo). In some cases, it is also useful to simplify the phase classification from supervised ML methods as a strictly binary outcome. For this purpose, we used a threshold of *OC* = 0.5 (i.e., a segment is classified as Ld if *OC* < 0.5 and Lo otherwise). The accuracy of binary classification was assessed from confusion matrices as the number of correct predictions divided by the total number of predictions. One notable advantage of binary classification is the ability to estimate domain size by counting runs of identical Lo or Ld segments within a vesicle.

## 3. Results and Discussion

### 3.1 Model system and generation of image datasets

The primary inputs for generating images of phase separated vesicles are the electron scattering profiles of Ld and Lo phases, which we calculated from MD simulations as described in Methods. We chose the three-component mixture DSPC/DOPC/Chol as a model system because the room temperature phase diagram has been determined experimentally at high compositional resolution (40,41). As shown in Fig. S1, the phase diagram contains a region of coexisting Ld and Lo phases that serve as a model for lipid rafts. We simulated the endpoints of a tieline that includes the composition DSPC/DOPC/Chol 39/39/22 mol%, in part because this composition has been extensively studied with confocal microscopy (42), FRET (43), X-ray scattering (24), and neutron scattering (3). The large line tension for this mixture (44) results in essentially complete coalescence of domains in giant unilamellar vesicles that can be easily visualized with fluorescence microscopy, such that each vesicle has a single domain of each phase.

Table 1 lists bilayer structural parameters of Ld and Lo phases obtained from the molecular simulations. Most important for cryo-EM are the substantial differences in bilayer thickness *D*_*s*_ and area per lipid *A*_*L*_, the latter being inversely related to lipid packing such that the smaller *A*_*L*_ of Lo phase indicates greater lipid packing density compared to Ld phase. The large differences in bilayer thickness and lipid packing result in markedly different electron scattering profiles for Ld and Lo bilayers, as shown in Fig. S2. We used these electron scattering profiles to generate image datasets as a function of Lo area percentage ranging from 0 to 100% in increments of 10%, as shown in Fig. 1, with the workflow for image generation demonstrated schematically in Fig. S2. Throughout the text, we refer to these compositions as S0-S10. For each Lo area percentage, 490 vesicle images were generated, with the size of individual vesicles randomly drawn from a distribution consistent with extrusion through a 50 nm pore size filter and thus producing diameters ranging from ∼ 20-80 nm. Each vesicle object was given a different random 3D orientation prior to projection. Projection images were convolved with a contrast transfer function (CTF) assuming an electron energy of 300 keV and using typical values for CTF parameters (Table 2). Figure 2 shows representative images from each simulated composition before and after the addition of Gaussian noise. For most of the phase separated vesicles (i.e., S1-S9), distinct domains of different thickness and intensity are visible in the noisy images.

**Figure 1.**
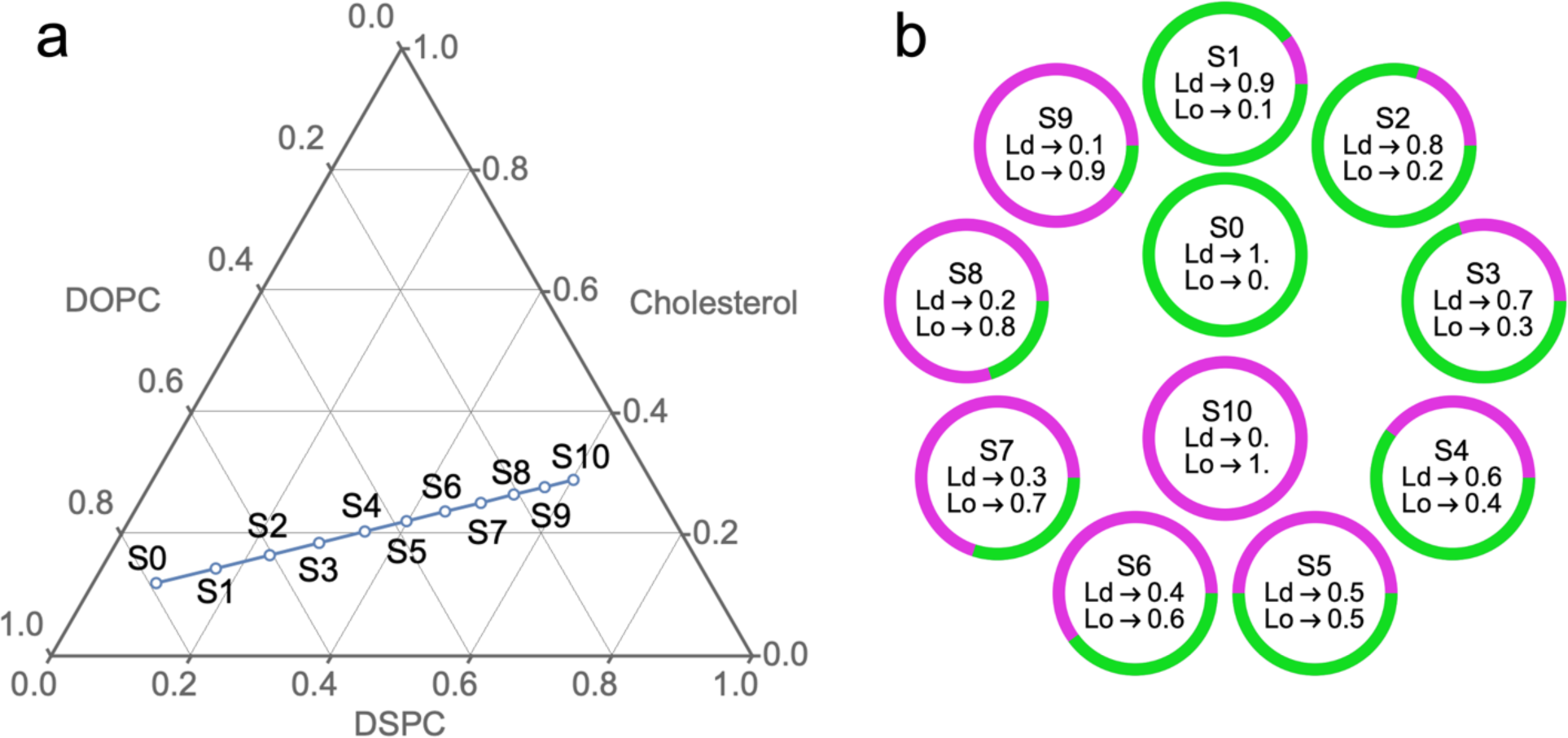
Lipid compositions of synthetic image datasets. (a) All compositions in this study lie on a thermodynamic tieline of the mixture DSPC/DOPC/Chol at 20°C (40). On this tieline, an Ld phase of fixed composition (S0) coexists with an Lo phase of fixed composition (S10) and only the relative amounts of Ld and Lo change. (b) The compositions S1-S9 represent a series of increasing Lo area percentage in increments of 10%.

**Figure 2.**
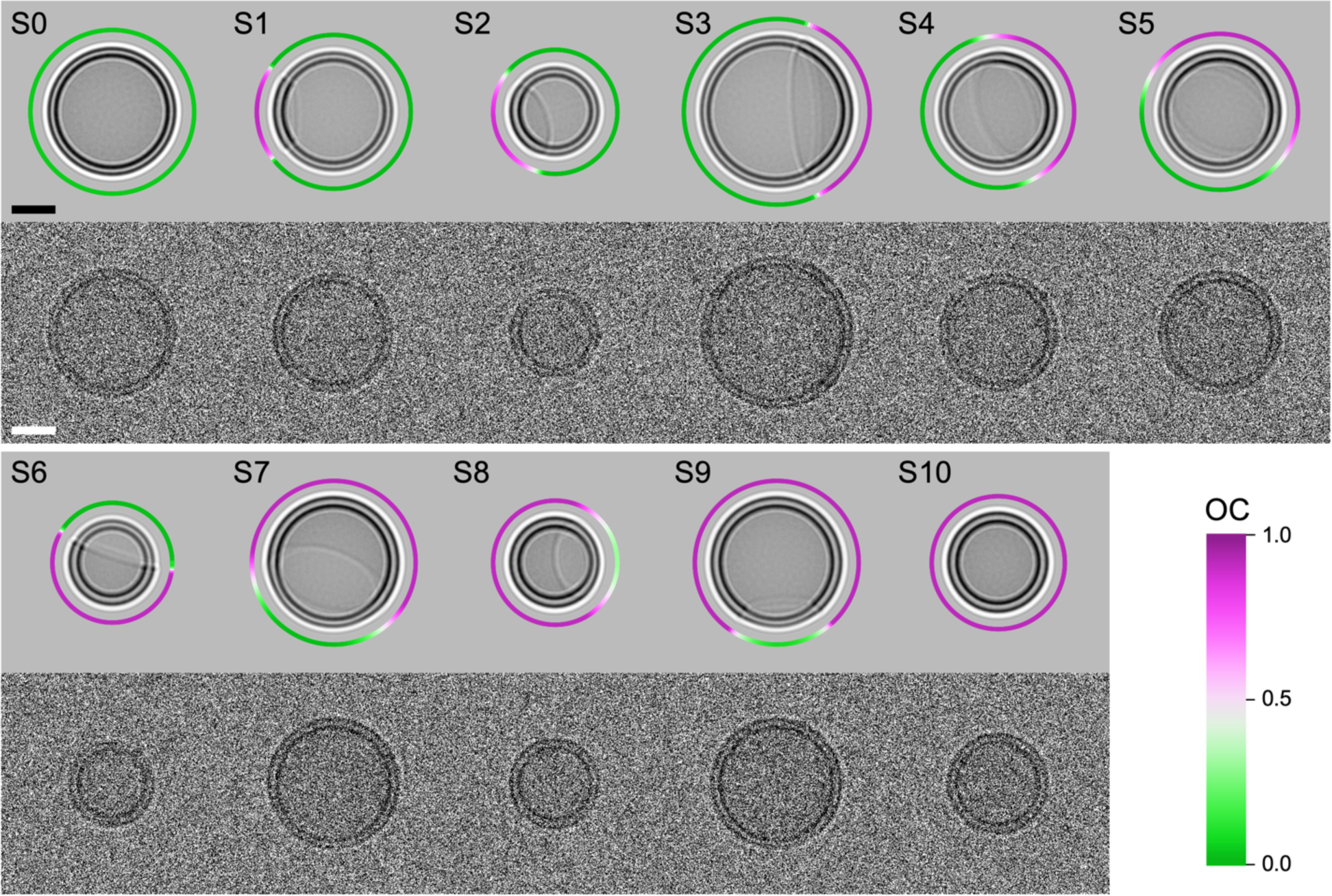
Representative vesicle images. A pair of images of a representative vesicle from each uniform (S0, S10) and phase separated (S1-S9) dataset is shown (see Fig. 1 for compositions). For each pair, the top and bottom images are before and after addition of Gaussian noise, respectively. The ring surrounding the vesicle in the noise-free images shows the angular dependence of the *OC* value in the projection, which quantifies the fraction of signal arising from the Lo domain. Scale bar 20 nm.

**Table 2.** Parameters used to generate simulated images (see Methods for details).

| *electron* |  | *contrast transfer function* |  |  |  |
| --- | --- | --- | --- | --- | --- |
| energy | $\lambda$ | $\Delta Z$ | $Q$ | $B$ | $m$ |
| 300 keV | 1.97 pm | 2.5 $\mu\text{m}$ | 0.075 | 300 $\text{\AA}^2$ | 400 |

Datasets of noisy images were then analyzed to obtain intensity profiles (IPs) around the vesicle circumference at 5 nm resolution as detailed in Methods. Figure 3 compares the ground truth profiles (obtained by convolving the Ld and Lo electron scattering profiles with the CTF) to representative IPs obtained from synthetic images of pure Ld and Lo vesicles. The ground truth IPs of Ld and Lo phases (Fig. 3b) are characterized by significant differences in both the position and depths of the “troughs” on either side of the bilayer center. These features, which arise from differences in bilayer thickness and molecular packing density of Ld and Lo phases, are partially obscured by noise at 5 nm resolution but are precisely recovered when IPs from many segments are averaged (Fig. 3c-d). As we now describe, the spatially resolved IPs constitute the raw data that we used to classify the bilayer phase state.

**Figure 3.**
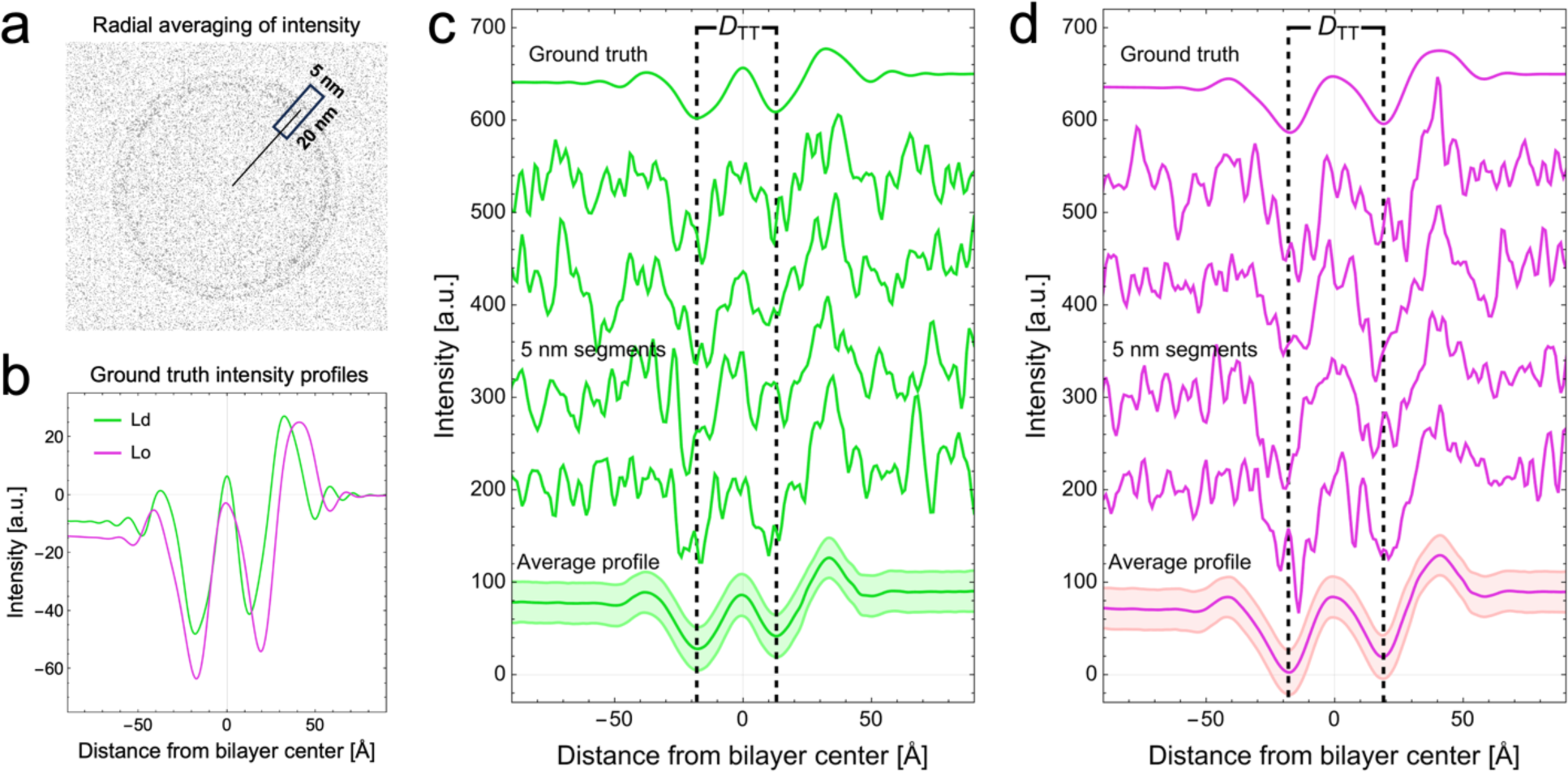
Segment intensity profiles. (a) Spatially resolved intensity profiles (IPs) are obtained by radial averaging of the intensity within segments of ∼ 5 nm arc length centered at the projected vesicle contour. (b) Ground truth IPs of Ld and Lo phases show significant differences in the position and depth of the troughs on either side of the bilayer center. (c and d) Comparison of Ld (c) and Lo (d) IPs: ground truth (top), four randomly chosen 5 nm segments (middle), and the average of all segments in all vesicles (bottom, with shaded area denoting ±1 standard deviation). IPs in panels c and d are offset vertically for clarity.

### 3.2 Phase classification based on bilayer thickness

As a baseline for comparing the performance of machine learning methods, we first employed a simple classification scheme for phase state based on bilayer thickness measured from segment IPs as described in Methods. Figure S3a shows probability distributions of segment thicknesses *D*_*TT*_ for each dataset. As expected, the distributions are approximately Gaussian for the single-phase Ld and Lo datasets S0 and S10, while datasets S1-S9 show non-Gaussian character owing to Ld+Lo coexistence within the vesicles. The mean *D*_*TT*_ was 29.7 Å for S0 and 35.8 Å for S10, closely matching the ground truth values of 30.6 Å and 36.4 Å for Ld and Lo phases, respectively.

To classify the phase state of individual segments in compositions S1-S9, we specified a cutoff thickness, 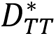, such that a segment whose thickness was greater than or equal to 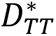 was classified as Lo. A confounding problem for this classification scheme is the broad and overlapping *D*_*TT*_ distributions of Ld and Lo phases, which precludes a straightforward determination of an appropriate cutoff. For example, using the midpoint between the mean Ld and Lo *D*_*TT*_ values (i.e., 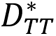 = 33 Å) resulted in an overestimation of the Lo percentage for samples S1-S6 and an underestimation for samples S8-S9 (Fig. S3b left). This discrepancy is a direct result of the narrower distribution of measured *D*_*TT*_ values for Lo compared to Ld segments. Using a larger value of 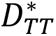 = 34 Å maximized the overall accuracy of phase classification at the level of individual segments (Fig. S3b middle, Table 4), achieving an average accuracy of 82% for all 11 systems. It is notable that no cutoff value was able to accurately predict phase area percentages for all tieline samples (Fig. S3b), a fact that severely limits the utility of this approach to phase classification.

### 3.3 Phase classification by unsupervised ML

We next used unsupervised learning (*k*-means and *k*-medoids) to group the segment IPs into two clusters (we note that for this analysis, segments from all 11 samples were pooled prior to clustering). As shown in Table 3, both clustering methods resulted in reasonable area percentage predictions for all compositions, deviating by < 5% from ground truth values. Both methods also achieved good accuracy (> 90%) at the level of individual segment classification (Table 4). The accuracy was correlated with the position of the sample on the tieline, with the lowest accuracy consistently occurring near the middle of the tieline (i.e., sample S5). To assess the impact of dimensionality reduction, we also performed clustering after first subjecting the IPs to a global principal component analysis. Figure S4 reveals that segment accuracy exceeds 90% for all datasets even when only the first two principal components are used for clustering. We note that for datasets of the size used here, *k*-means and *k*-medoids clustering are computationally inexpensive and fast on a desktop computer even when the full 201-dimensional IP is used.

**Table 3.** Area percentage of Lo phase determined by different phase classification methods.

| Sample | Ground truth | *Thickness* | *Unsupervised ML* |  | *Supervised ML* |  |  |  |  |
| --- | --- | --- | --- | --- | --- | --- | --- | --- | --- |
| | | $D_{TT}^* = 34 \text{ \AA}$ | 2-means cluster | 2-medoids cluster | DT | GBT | RF | LR | NN |
| S0 | 0.0 | 15 | 2 | 2 | 6 | 2 | 2 | 1 | 1 |
| S1 | 10.0 | 23 | 11 | 10 | 16 | 12 | 11 | 12 | 11 |
| S2 | 20.0 | 30 | 20 | 20 | 25 | 22 | 21 | 21 | 21 |
| S3 | 30.0 | 37 | 31 | 30 | 34 | 32 | 31 | 32 | 31 |
| S4 | 40.0 | 44 | 39 | 38 | 42 | 41 | 40 | 41 | 40 |
| S5 | 50.0 | 50 | 48 | 48 | 50 | 50 | 49 | 50 | 49 |
| S6 | 60.0 | 57 | 58 | 57 | 59 | 60 | 58 | 60 | 59 |
| S7 | 70.0 | 64 | 68 | 67 | 69 | 70 | 69 | 71 | 69 |
| S8 | 80.0 | 70 | 77 | 76 | 77 | 79 | 77 | 79 | 78 |
| S9 | 90.0 | 77 | 86 | 85 | 85 | 88 | 87 | 89 | 88 |
| S10 | 100.0 | 84 | 96 | 96 | 93 | 98 | 97 | 99 | 97 |

**Table 4.** Segment-level accuracy determined by different phase classification methods.

| Sample | *Thickness* | *Unsupervised ML* |  | *Supervised ML* |  |  |  |  |
| --- | --- | --- | --- | --- | --- | --- | --- | --- |
| | $D_{TT}^* = 34 \text{ \AA}$ | 2-means cluster | 2-medoids cluster | DT | GBT | RF | LR | NN |
| S0 | 84 | 98 | 98 | 94 | 98 | 98 | 99 | 99 |
| S1 | 82 | 94 | 94 | 91 | 95 | 95 | 95 | 95 |
| S2 | 82 | 93 | 93 | 89 | 94 | 94 | 94 | 94 |
| S3 | 81 | 93 | 92 | 89 | 94 | 94 | 94 | 94 |
| S4 | 82 | 92 | 92 | 89 | 94 | 93 | 94 | 93 |
| S5 | 81 | 91 | 91 | 88 | 93 | 92 | 93 | 92 |
| S6 | 81 | 92 | 91 | 88 | 93 | 92 | 93 | 93 |
| S7 | 82 | 93 | 92 | 90 | 94 | 93 | 95 | 94 |
| S8 | 82 | 92 | 92 | 90 | 94 | 93 | 95 | 94 |
| S9 | 82 | 93 | 92 | 90 | 95 | 93 | 95 | 94 |
| S10 | 84 | 96 | 96 | 93 | 98 | 97 | 98 | 98 |

### 3.4 Phase classification by supervised ML

We next tested five supervised ML algorithms as described in Methods. Models were trained on segment IPs from the Ld and Lo datasets, with 30% of these segments reserved for post-training accuracy assessment. Table 3 compares the Lo area percentage predicted by these models to predictions from thickness-based and unsupervised methods, while Table 4 compares the overall accuracy of binary phase classification for the various datasets and methods at the level of individual segments. Like the unsupervised methods, accuracy is strongly correlated with the position of the sample on the tieline (Fig. 4a). Comparing different methods, Gradient Boosted Trees (GBT) and Logistic Regression (LR) emerged as the most accurate overall, followed closely by Nearest Neighbors (NN) and Random Forest (RF): Decision Tree (DT) was significantly less accurate. Figure S5 shows an example of the confusion matrices that were used to calculate binary accuracy.

**Figure 4.**
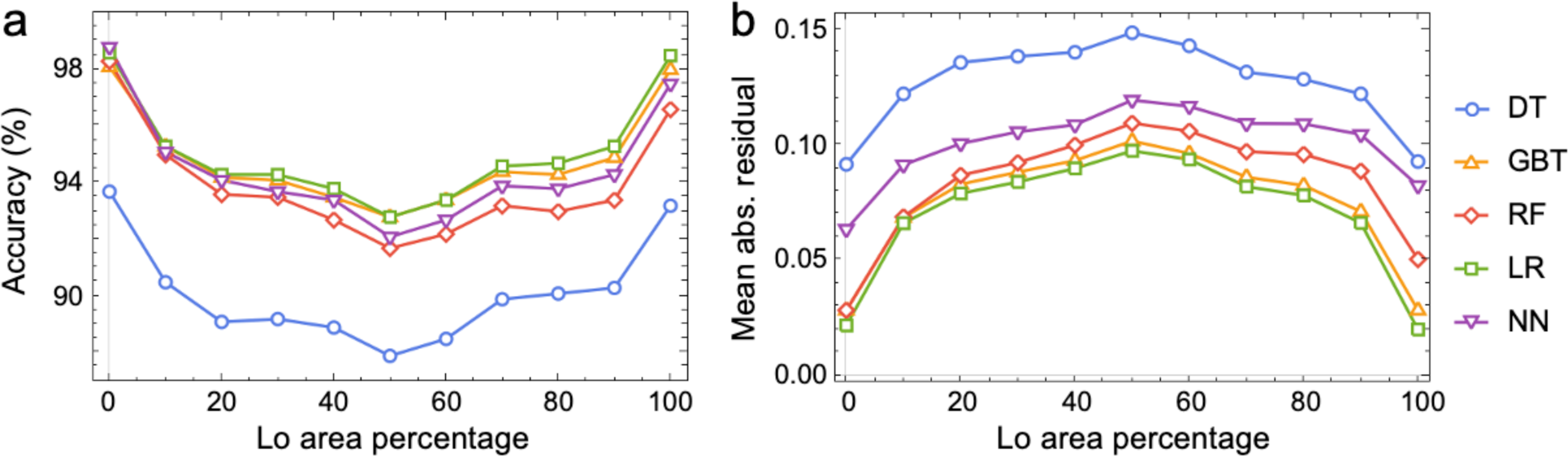
Accuracy of phase classification by supervised ML methods. (a) Percentage of correctly predicted segments vs. Lo area percentage. (b) Mean absolute residual of segment ordered character as defined in the text. Methods used: Decision Tree (DT), Gradient Boosted Trees (GBT), Random Forest (RF), Logistic Regression (LR), and Nearest Neighbors (NN).

Because supervised methods yield probability-based predictions (i.e., the probability that a given segment is Lo phase), we can further assess the ability of these models to recognize domain boundaries that are inherently weighted superpositions of Lo and Ld IPs. For each segment, we first calculated the fraction of total intensity originating from the Lo phase, which we termed the segment’s ordered character (*OC*). The residual *OC* − *OC*′, where *OC* is the ground truth value and *OC*′ is the probability determined by a supervised model, is a natural alternative metric for quantifying accuracy, with smaller values indicating greater accuracy. The mean absolute residual (i.e., averaged over all segments in a dataset) is plotted vs. Lo area percentage in Fig. 4b for the different supervised methods, revealing similar overall trends compared to the binary phase classification. The accuracy of boundary classification is discussed in the next section.

### 3.5 Visualizing phase classification accuracy

It is instructive to visualize the accuracy of phase classification at the vesicle level. Figure 5 presents ring plots of representative vesicles in which ground truth *OC* values of individual segments are found in the innermost ring and *OC*′ values predicted by the various supervised models are arranged concentrically. The green-pink color scale facilitates the identification of misclassified segments (i.e., an Lo segment identified as Ld or vice versa), in this case when a binary cutoff of *OC*^∗^ = 0.5 was used, as described in Methods. A few such misclassified segments are found in each of the phase separated vesicles shown in Fig. 5. Unsurprisingly, the probability of misclassification is greater near domain boundaries where both phases contribute to the projected signal. Notably, the representative examples of pure Ld and Lo vesicles shown in Fig. 5b and 5g contain no misclassifications, consistent with the absence of confounding boundary segments in these datasets, although misclassifications in the bulk do occasionally occur in both uniform and phase-separated vesicles and can be seen in Fig. 5c, 5e and 5f.

**Figure 5.**
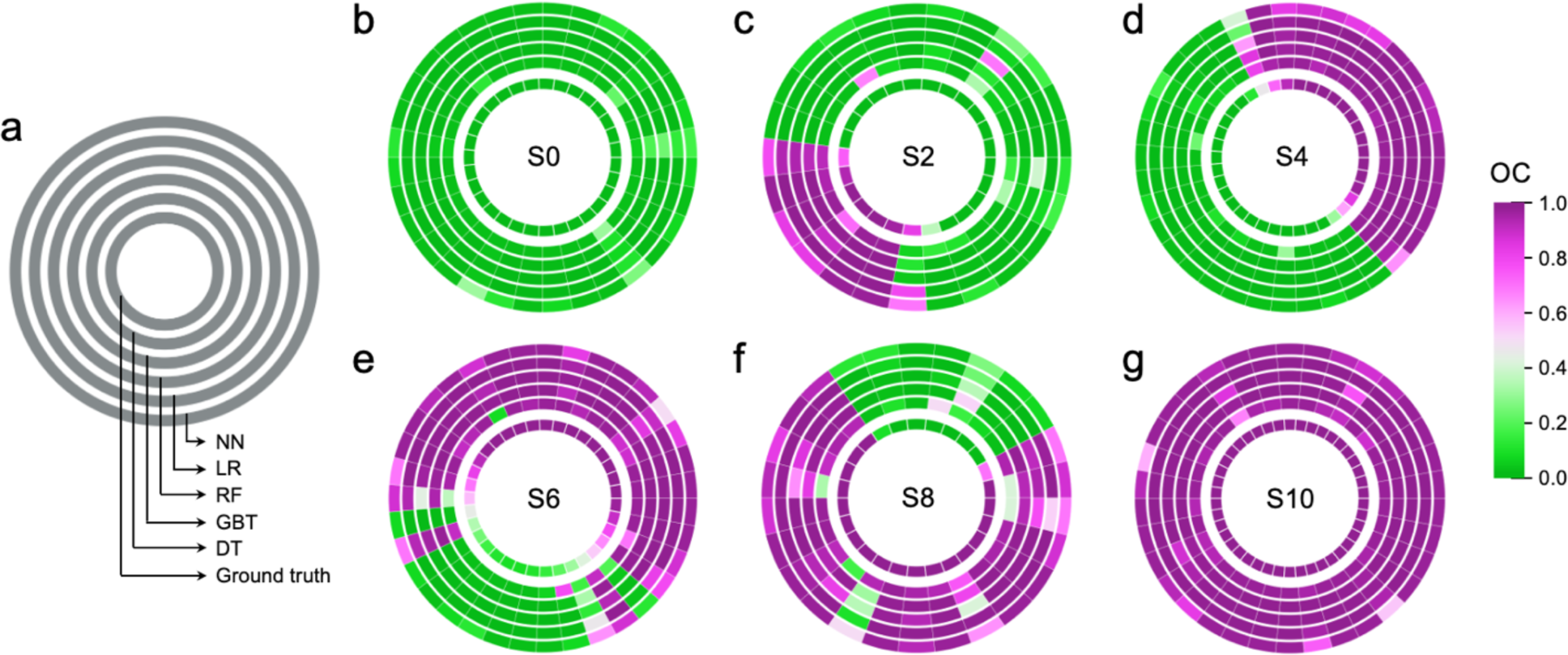
Visualizing the accuracy of phase classification for individual vesicles. (a) Legend: the inner ring gives the ground truth *OC* for individual 5 nm segments of a vesicle, with model predictions *OC*′ arranged concentrically as indicated. (b-g) Ring plots of representative vesicles for different datasets as indicated. Methods used: Decision Tree (DT), Gradient Boosted Trees (GBT), Random Forest (RF), Logistic Regression (LR), and Nearest Neighbors (NN).

In light of the previous discussion, the trends in accuracy shown in Fig. 4 can be explained by the difficulty of classifying boundary segments. The lowest overall accuracy occurs in system S5, which has equal area fractions of the two phases and thus the largest number of boundary segments. Indeed, every projection image for this sample—regardless of the vesicle’s 3D orientation—is assured to have boundary segments, as demonstrated in Fig. 6a-f. However, as the vesicle composition moves toward the ends of the tieline in either direction, the size of one domain increases at the expense of the other, and the total domain boundary length in the projected images decreases. Consequently, there is a non-zero probability of observing only the majority phase at the edges of a projected vesicle, as demonstrated in Fig. 6l for sample S2. Such projections are relatively common for compositions near the ends of the tieline (Fig. S6), and as a result, the accuracy of phase identification is higher than for samples near the middle of the tieline.

**Figure 6.**
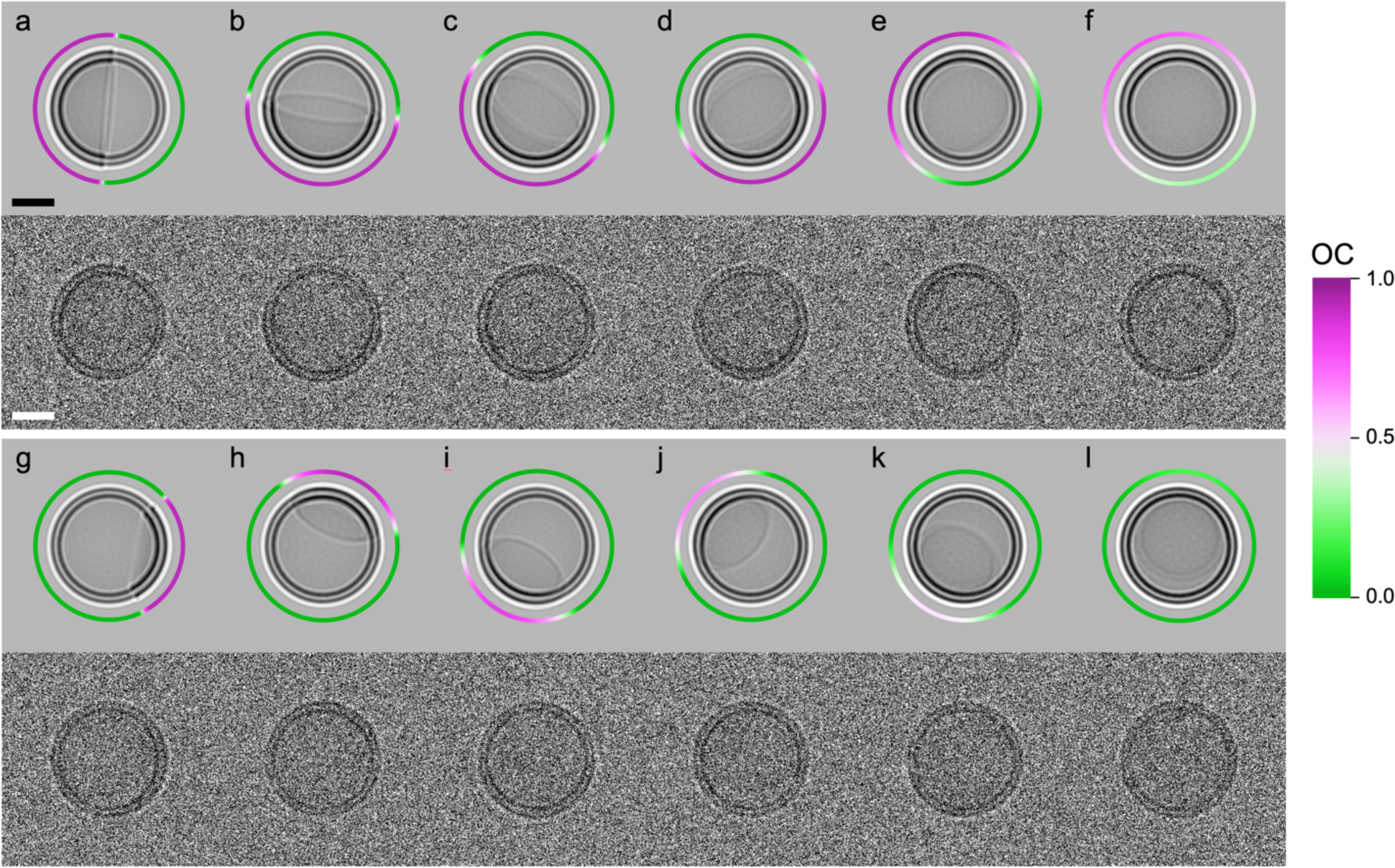
Selected images demonstrating the influence of vesicle orientation on apparent phase behavior. Show are pairs of images for several orientations of ∼ 50 nm diameter vesicles from datasets S5 (a-f) and S2 (g-l). In each pair, the upper and lower images are before and after the addition of Gaussian noise, respectively. The ring surrounding the vesicle in the noise-free images plots the ground truth *OC* value along the projected circumference as indicated by the color scale. Scale bar 20 nm.

Further insight can be gained by visualizing segment classification accuracy for an entire dataset. For this purpose, we generated matrix plots in which rows correspond to individual vesicles (arranged from smallest to largest diameter, top to bottom) and columns correspond to individual segments (arranged such that the middle of the minority-phase domain is centered within the row), as shown in Fig. 7 for system S4. Several interesting aspects of the datasets become apparent in these plots. For example, the ground truth *OC* values plotted in Fig. 7a explicitly show how the fraction of boundary segments (i.e. those with lighter shades) varies widely between vesicles due to differences in their random 3D orientations prior to projection (cf. Fig. 6). In a few cases, the vesicle was oriented such that the domain boundary was nearly parallel to the projection plane, resulting in every segment containing intensity contributions from both phases. Figure 7b plots the corresponding *OC*′ values predicted by GBT, revealing a strong tendency for the model to predict values close to 0 or 1 and eschew intermediate values, even for boundary segments whose ground truth *OC* values are in fact intermediate. This tendency is emphasized in Fig. 7c, which plots the signed residual *OC* − *OC*′ and thus makes clear the outsized contribution of boundary segments to overall accuracy. The relatively poor performance in boundary identification is likely caused by the fact that the models were not trained to recognize boundary segments. Adding such segments to the training dataset will almost certainly improve the predictions and may prove to be important for real-world data analysis, where the likelihood of encountering multiple smaller domains within vesicles is greater.

**Figure 7.**
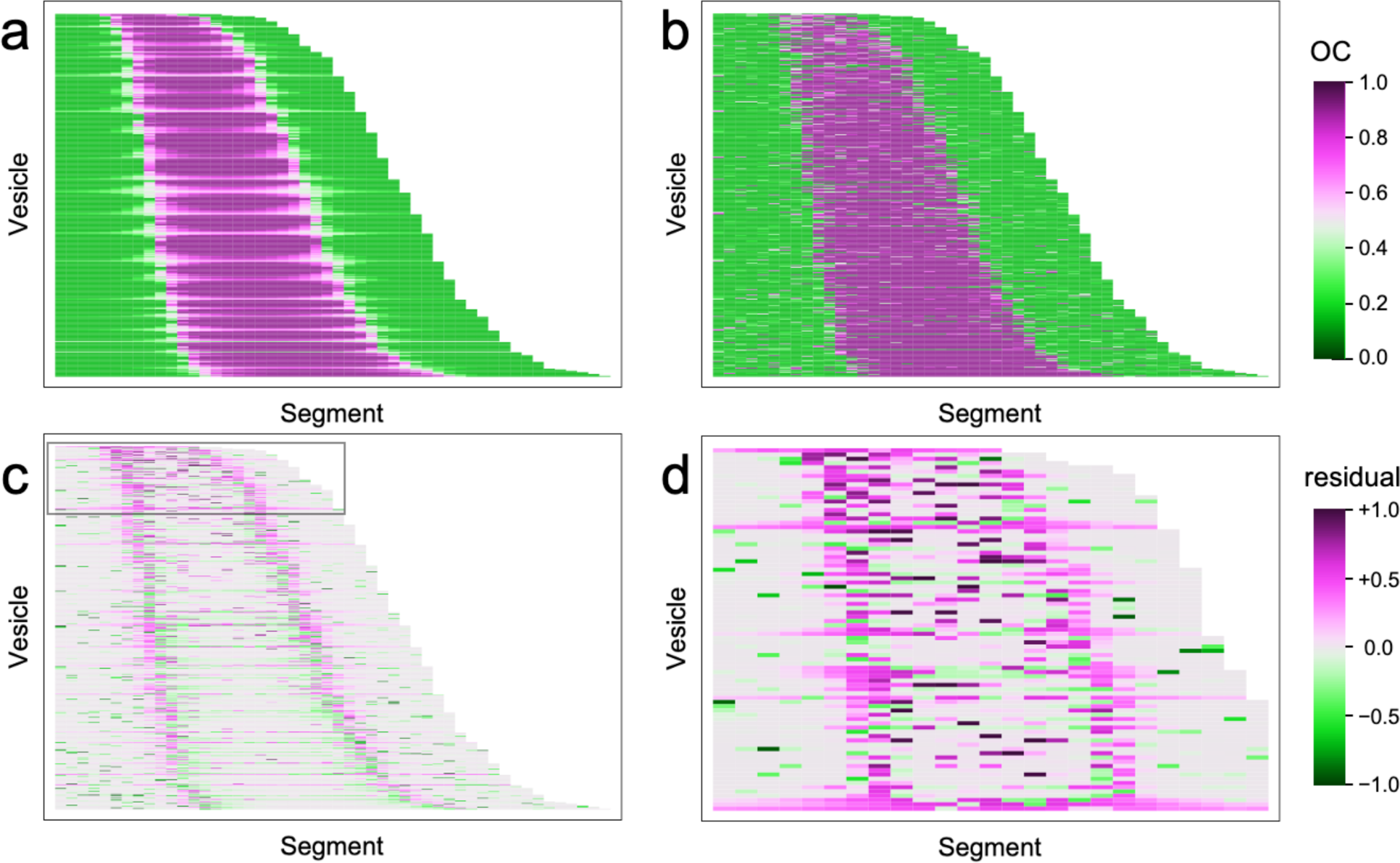
Matrix plots of ordered character. Each row represents one of 490 vesicles in dataset S4, where the columns are individual 5 nm segments of the vesicle. Plotted are: (a) the ground truth *OC* value; (b) the *OC*′ prediction from the GBT model; (c) the signed residual *OC* − *OC*′. Panel (d) expands the boxed region in panel (c) containing the subset of vesicles with diameters < 40 nm.

Figure 7 also sheds light on the relationship between vesicle size and accuracy of phase classification. This is most clearly seen in Fig. 7d, which expands the boxed region of Fig. 7c to highlight that segment misclassification is exacerbated in vesicles with diameters < 40 nm. There are likely two factors contributing to this phenomenon. First, smaller vesicles possess an inherently greater fraction of boundary segments on average. A second factor is the reduction in total signal that occurs when electrons encounter less lipid material on their path to the detector, as is the case for smaller vesicles. This effect is demonstrated by the nearly two-fold increase in contrast with increasing vesicle size shown in Fig. 8, as well as by Fig. S7, which plots the contrast of individual vesicles (defined as the intensity difference between the outer peak and inner trough) for the Ld and Lo samples. As shown in Fig. S7b, the contrast from an Lo segment in a very small vesicle is comparable to the contrast of an Ld segment from a vesicle near the peak of the size distribution, suggesting that it is more likely for an Lo segment from a small vesicle to be misclassified as Ld than vice versa. Indeed, this is precisely what is found for smaller vesicles as evidenced by the more frequent occurrence of pink shading (i.e., positive *OC* residuals, indicating an Lo segment misclassified as Ld) compared to green shading.

**Figure 8.**
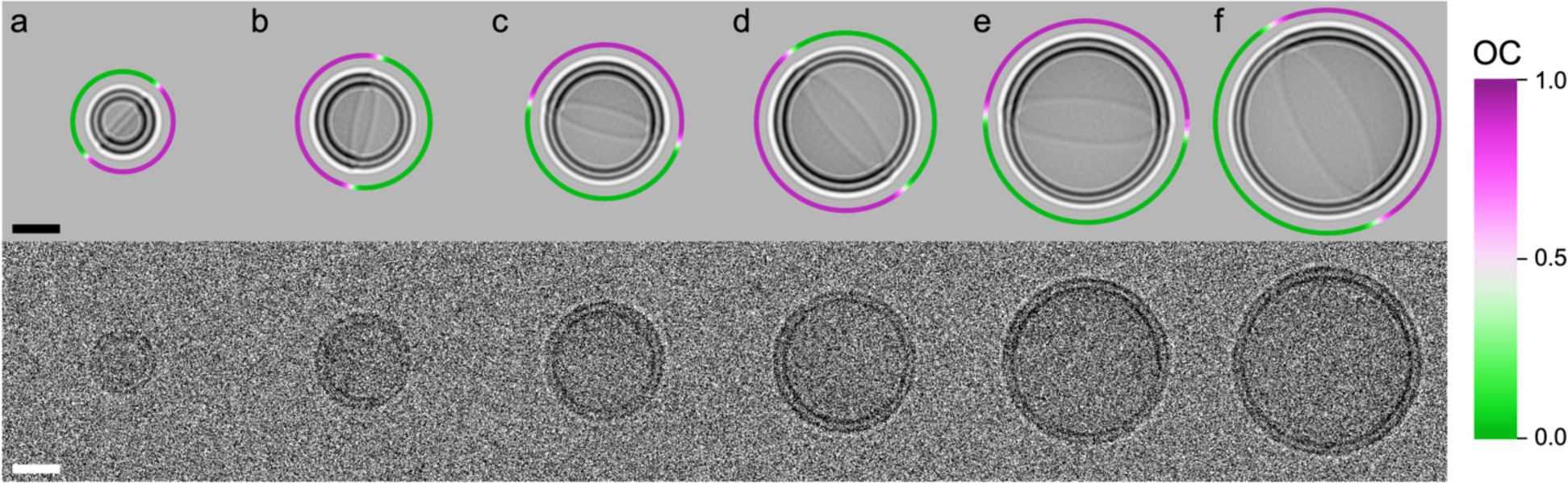
Vesicle size influences contrast. Shown are pairs of images of selected vesicles from dataset S5. In each pair, the upper and lower images are before and after the addition of Gaussian noise, respectively. The ring surrounding the vesicle in the noise-free images plots the ground truth *OC* value along the projected circumference as indicated by the color scale. Scale bar 20 nm.

### 3.6 Average IPs of coexisting phases

Among the most important information obtained from cryo-EM analysis is the average IP of the bilayer. In principle, the bilayer’s electron scattering profile can be recovered from the IP by deconvolution (45), raising the possibility of using cryo-EM to complement well-established neutron and X-ray based methods for bilayer structure determination (46). A significant advantage of cryo-EM compared to traditional scattering methods is the ability to classify individual segments by their phase and thus separately determine the IPs of coexisting phases. Figure 9a compares the mean IPs for Ld and Lo phases (i.e., averaged over all segments classified as Ld or Lo) for samples S0-S10 using predictions from GBT. As expected, agreement with ground truth profiles (Fig. 3) is better for the majority phase in each dataset, although the overall agreement for both phases is reasonable in all datasets. Figure 9b shows that *D*_*TT*_ values calculated from the IPs of phase separated samples S1-S9 are generally in excellent agreement with the values calculated from the pure Ld and Lo datasets (dashed lines) although deviations occur for the minority phase near tieline endpoints.

**Figure 9.**
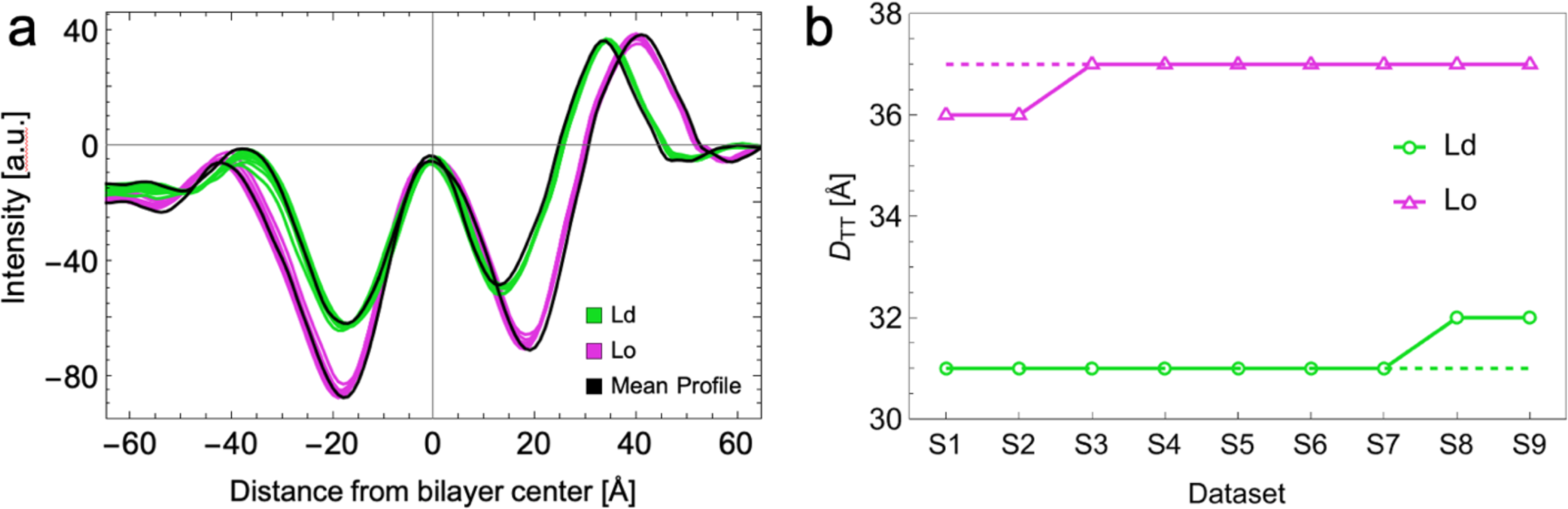
Average intensity profiles and *D*_*TT*_ values. (a) After classification by GBT, the mean intensity profile for all Lo (Ld) segments is depicted in pink (green) for each phase separated dataset (i.e., S1-S9), while the mean intensity profiles of pure Ld and Lo (datasets S0 and S10, respectively) are shown in black. (b) *D_TT_* values evaluated from the mean intensity profiles shown in (a). Dashed pink and green lines represent the *D_TT_* value of the pure Ld and Lo datasets.

### 3.7 Isolating vesicle size effects

It is well established that nanoscopic membrane curvature plays a vital role in many biological processes (47,48). Curvature is influenced by intrinsic factors such as lipid composition and the presence of integral membrane proteins, as well as by external factors including cytoskeletal attachments and the binding of peripheral membrane proteins (49). Even in simple liposomes with nominally identical leaflet compositions, it is likely that substantial structural perturbations emerge in the presence of high membrane curvature. This has been observed in coarse-grained molecular dynamics simulations of 20 nm diameter vesicles, which showed changes in the headgroup and acyl chain packing of inner and outer leaflets that depended both on lipid headgroup type and tail saturation (50). In this context, another major advantage of cryo-EM compared to neutron and X-ray scattering is the ability to isolate membranes of different average curvature within the sample and separately interrogate their structure.

As a proof-of-principle demonstration, we binned vesicles in each dataset by size and calculated several properties for the different bins (we note that all vesicles in our simulated datasets have identical bilayer structure irrespective of their size). Figure 10 shows the apparent phase area fractions for systems S1-S9 calculated separately for each vesicle size bin. Consistent with our interpretation of Fig. 7d, there is a selective tendency for Lo segments to be misclassified as Ld in smaller vesicles which manifests as a systematic overestimate (underestimate) of the average Ld (Lo) phase fraction (Fig. 10) for vesicles smaller than 40 nm in diameter. One possibility for eliminating this artifact is to employ different models that are specifically trained for vesicles in a given size range.

**Figure 10.**
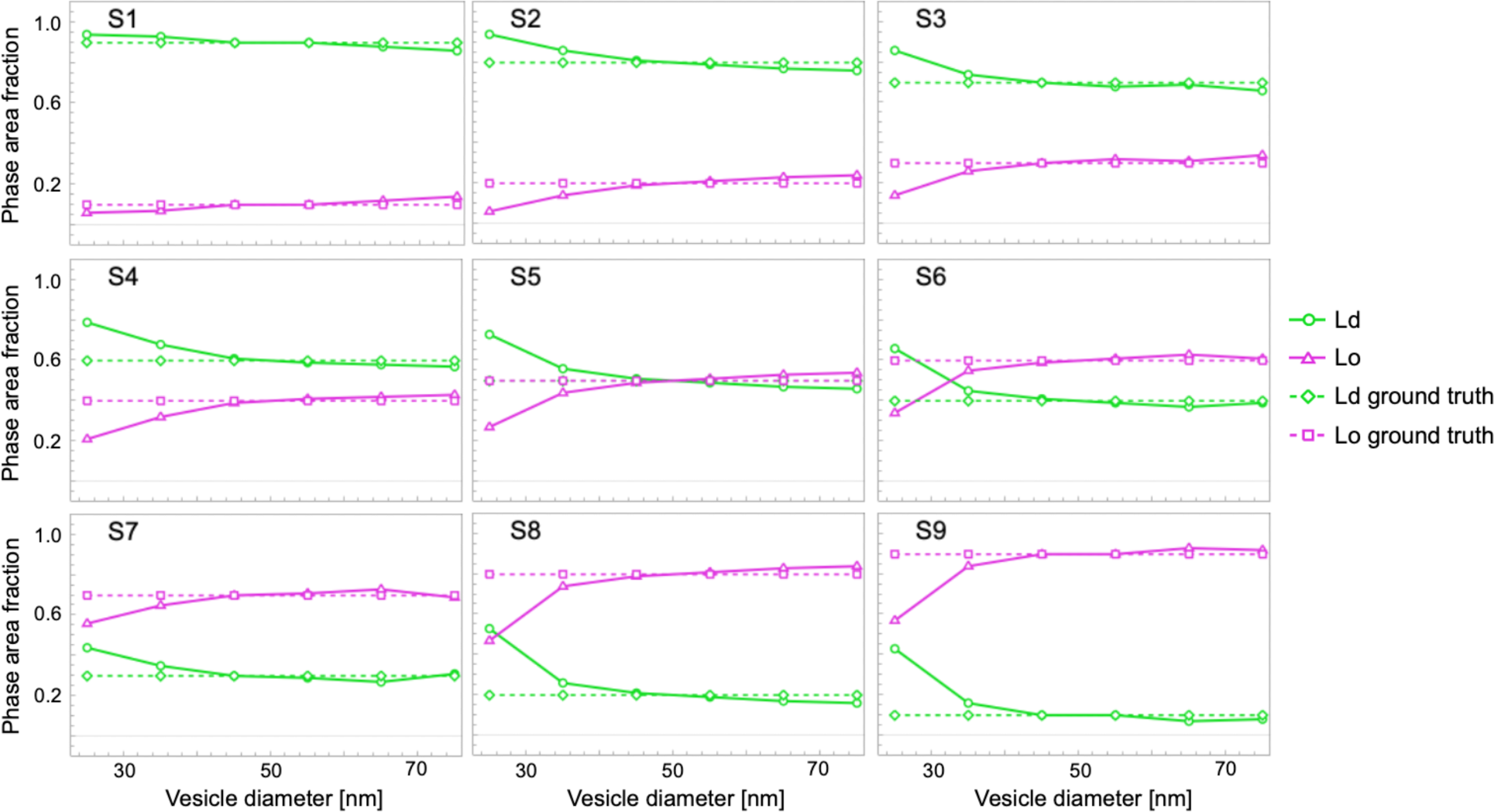
Influence of vesicle size on apparent phase area fractions. The area fraction of Ld (green) and Lo (pink) phases predicted by LR is shown as a function of vesicle size for each of the phase separated datasets. In all cases, there is a systematic overestimate of the Ld fraction for vesicles smaller than 40 nm diameter.

We also attempted to determine the average linear extent and number of domains as a function of vesicle size. This was complicated by the non-negligible frequency of segment misclassifications far from domain boundaries (cf. Fig. 5), which have the effect of erroneously dividing a continuous domain into two or more discontinuous domains. To counteract this artifact, we implemented a correction that eliminated isolated segments (i.e., those whose phase classification was different than its two nearest neighbors) as illustrated in Fig. S8a. With this correction, the average domain size calculated from images was similar to ground truth values (Fig. 11). Reasonably good agreement between the average number of domains per vesicle calculated from images and the ground truth value (i.e., 2 domains per vesicle) was also obtained from this analysis (Fig. S8b-c).

**Figure 11.**
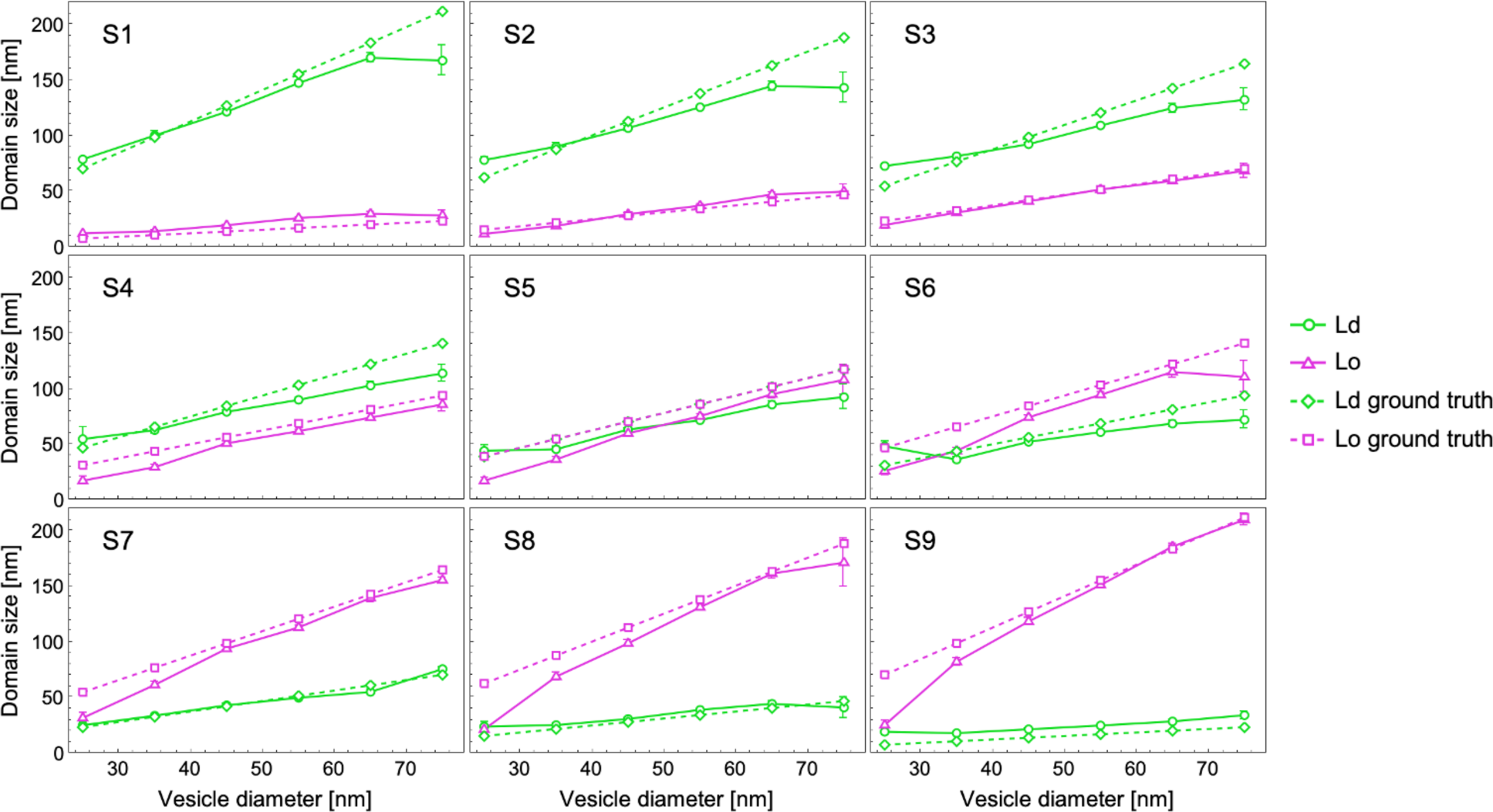
Influence of vesicle size on apparent domain size. The computed domain size across all nine systems is presented to examine the impact of vesicle sizes on area fractions. Logistic Regression (LR) is employed as the classification technique. Smaller vesicles often deviate from the ground truth values represented by dashed lines in their respective phases.

## 4. Summary and Conclusions

We report on a new application of machine learning to classify the phase state of lipid bilayers with 5 nm lateral resolution in cryo-EM images. To mimic the scenario of lipid rafts in eukaryotic plasma membranes, we analyzed synthetic images of phase separated vesicles constructed from MD simulations of Ld and Lo phases. The use of ground truth datasets allowed us to compare the accuracy of different methods, in addition to testing the performance of the models in the presence of heterogeneities in both vesicle size and domain size.

Our most important finding is that ML methods can classify the phase state of the bilayer at accuracies exceeding 90% for the conditions that were simulated. Supervised methods such as Logistic Regression and Gradient Boosted Trees showed the highest accuracy, while unsupervised methods such as *k*-means and *k*-medoids closely followed. Notably, unsupervised methods exhibited greater accuracy than some supervised methods, showcasing their potential for analyzing standalone experimental datasets and characterizing nanodomains in the absence of training data. In comparing results from supervised and unsupervised ML methods, we do not wish to imply that one is intrinsically superior to another; indeed, compared to classification based on segment thickness alone, each of the ML methods showed significantly improved performance. Instead, the choice between methods depends on the scientific questions being asked and the data that are available.

In cases where the models failed, some common trends emerged: (1) accuracy decreased for samples positioned in the middle of a tieline where approximately equal fractions of each phase are present; (2) accuracy decreased for vesicles smaller than ∼ 40 nm diameter. Each of these trends is ultimately related to the presence of domain boundaries that are particularly prone to misidentification in a binary classification scheme. Extensive domain interface is a hallmark of nanoscopic phase separation, and the ability to correctly classify boundary segments is thus a key target for future development. It must also be recognized that additional challenges are encountered *in vitro* and *in vivo* that are not accounted for in our *in silico* datasets; these include variability in physical parameters such as vesicle composition and ice thickness, as well as instrumental parameters like defocus length (26). Despite these limitations, the ability to directly image and decode nanoscopic membrane features will likely position cryo-EM at the leading edge of membrane biophysics in the years to come.

## Author contributions

K.S., M.D., M.N.W., and F.A.H. designed the research. M.D. performed molecular simulations. F.A.H. generated the synthetic datasets. K.S. and F.A.H. analyzed the datasets. K.S., M.D., M.N.W., and F.A.H. wrote the manuscript.

## Acknowledgements

We are grateful to Dr. Tristan Bepler for providing the machine learning algorithm, MEMNET, for automated contouring of vesicles. This work was supported by NSF Grant CHE-2204126 (to F.A.H and M.N.W.). M.N.W. acknowledges the William Wheless III Professorship.

