## Supporting Information for "Cryo-EM images of phase separated lipid bilayer vesicles analyzed with a machine learning approach"

**This PDF file includes:**

Figures S1 to S8
Supporting References

### SI Figures

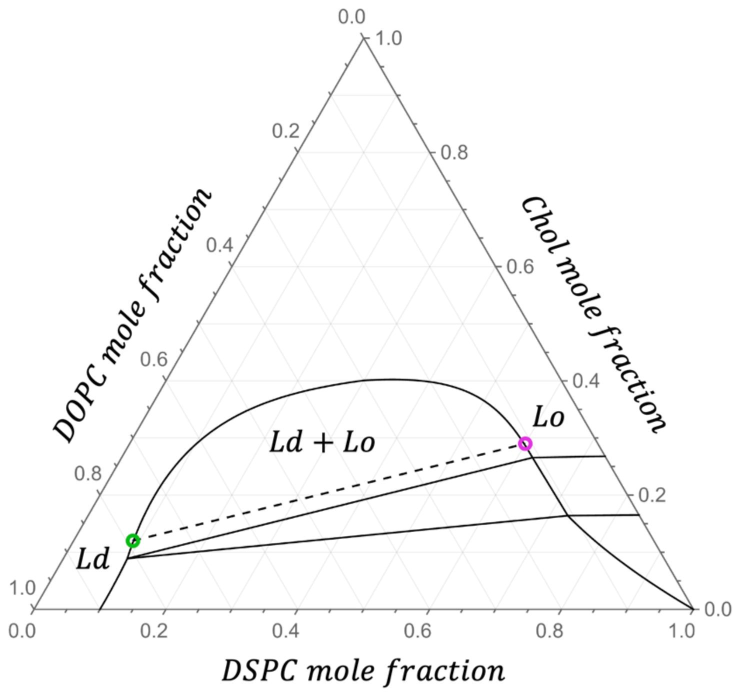


**Figure S1 Experimental phase diagram used in this study.** The 20°C phase diagram of DSPC/DOPC/Chol (1,2) showing the Ld+Lo coexistence region and the Ld and Lo endpoints (green and magenta circles, respectively) of the tieline that passes through the composition $\chi_{DSPC} = 0.39$, $\chi_{DOPC} = 0.39$, $\chi_{Chol} = 0.22$.

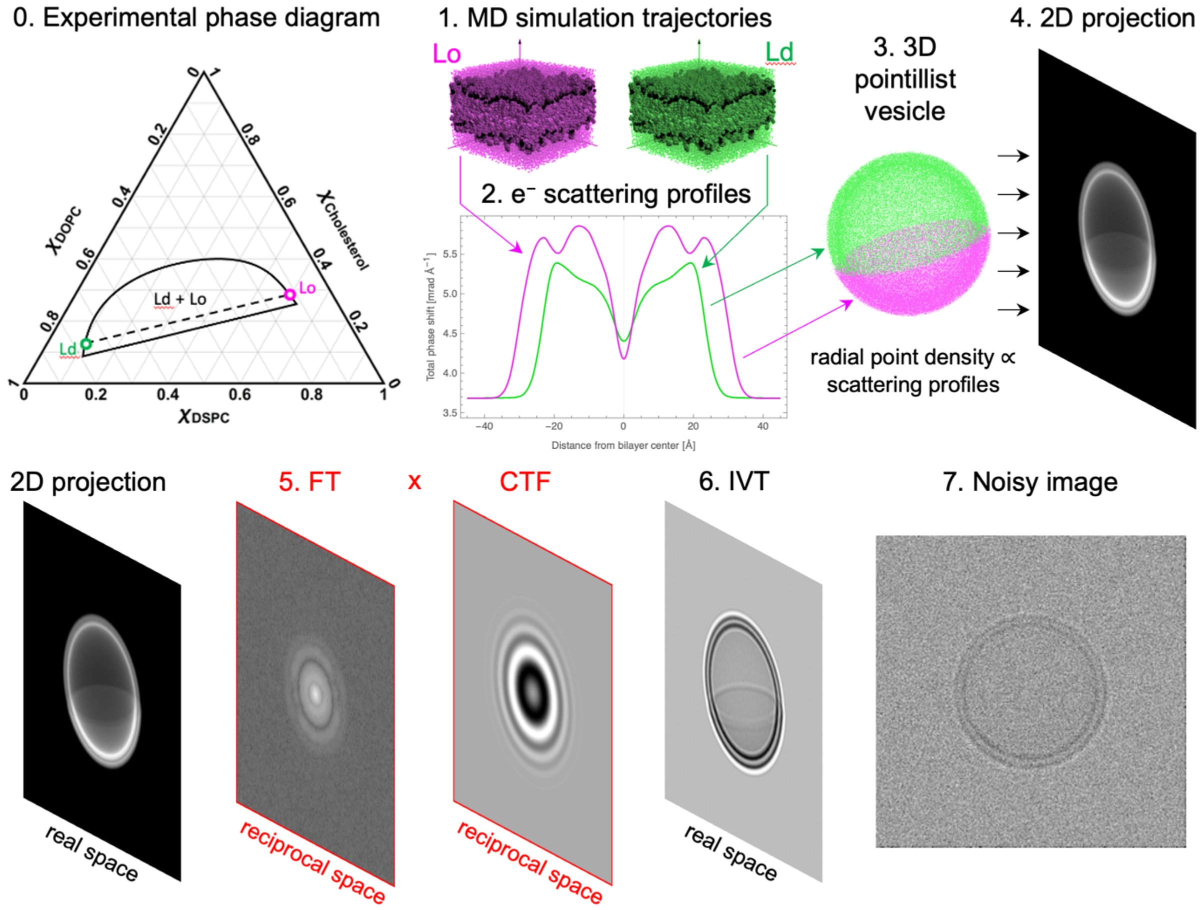


**Figure S2 Schematic of the workflow for generating synthetic cryo-EM images of lipid bilayer vesicles.** Step 0: The experimental phase diagram showing the Ld and Lo compositions (green and magenta circles, respectively). Steps 1 and 2: MD simulation trajectories are used to generate electron scattering profiles. Step 3: A 3D model of a phase separated vesicle is generated in which the density of points in each domain is proportional to the scattering profile of the respective phases. Step 4: the vesicle is rotated randomly about its origin and the points are projected onto a plane. Steps 5-6: the projected image is smeared by the instrument's contrast transfer function. Step 7: random Gaussian noise is added to each pixel.

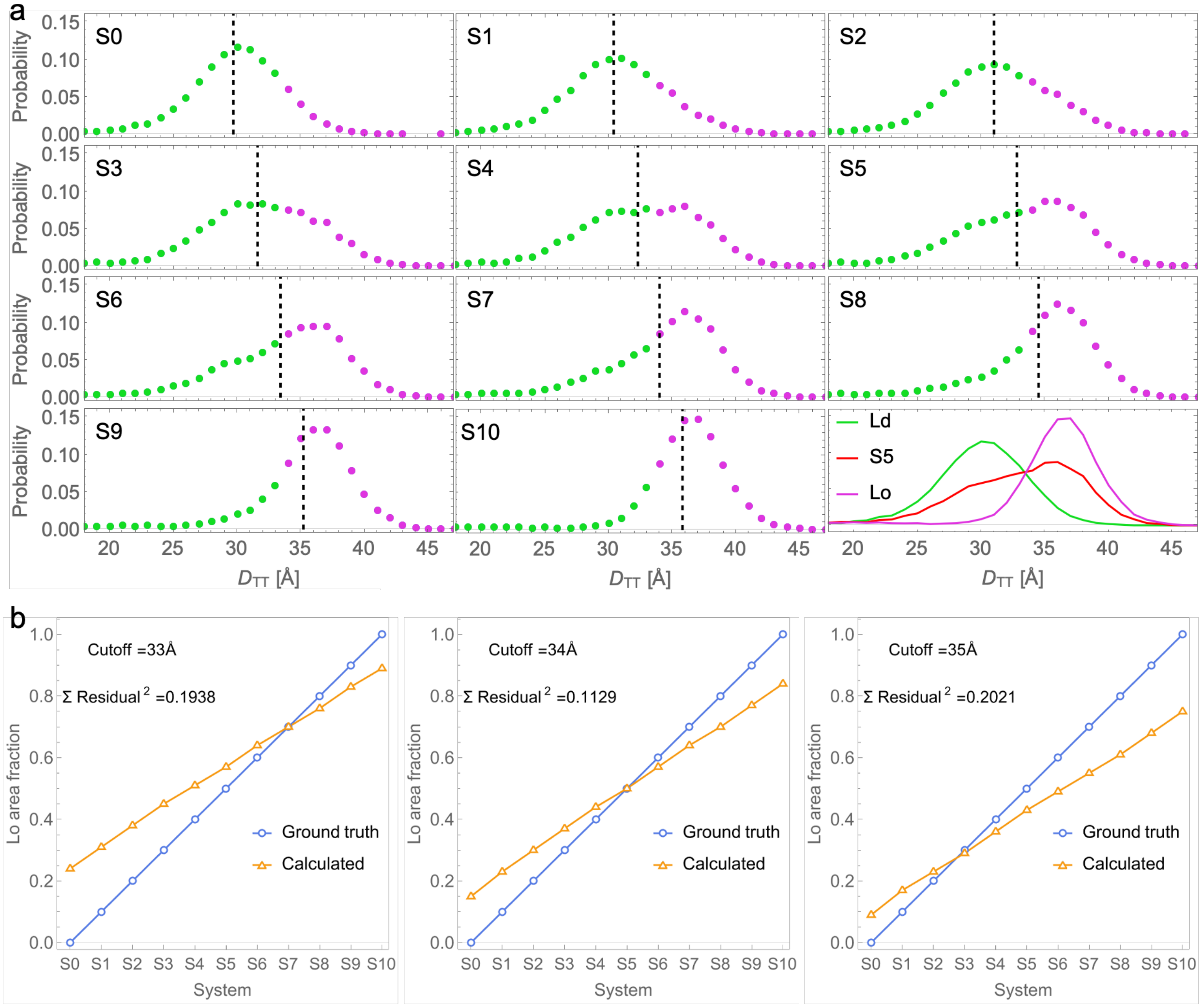


**Figure S3 Phase classification by segment thickness.** (a) Probability distribution of segment $D_{TT}$ values for datasets S0-S10. Segments with $D_{TT}$ values less than 34 Å are shown in green, while those exceeding this threshold are shown in pink. The mean segment $D_{TT}$ value is represented by a dashed line, providing a visual reference for the systematic increase in $D_{TT}$ with increasing fraction of Lo phase. (b) Predicted Lo area fraction (orange symbols) using different thickness cutoffs for segment classification as indicated on the figure.

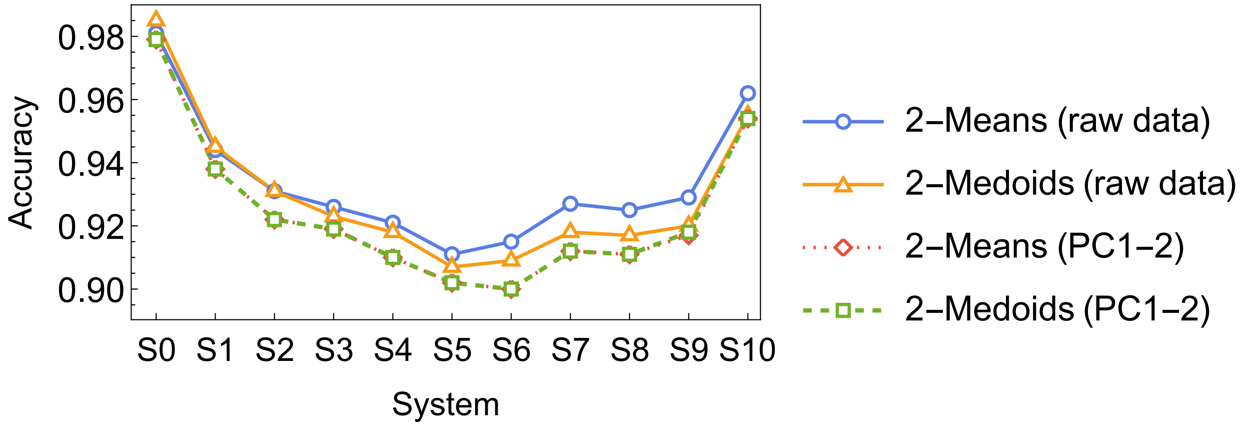


**Figure S4 Phase classification accuracy of unsupervised ML methods.** Shown is the segment-level accuracy for datasets S0-S10 from *k*-means (blue) or *k*-medoids (orange) clustering using either the full IP (solid shading) or the first two principal components (hatched shading). When considering the first two principal components, the centroids for 2-means and medoids for 2-medoids were found to be in close proximity to each other.

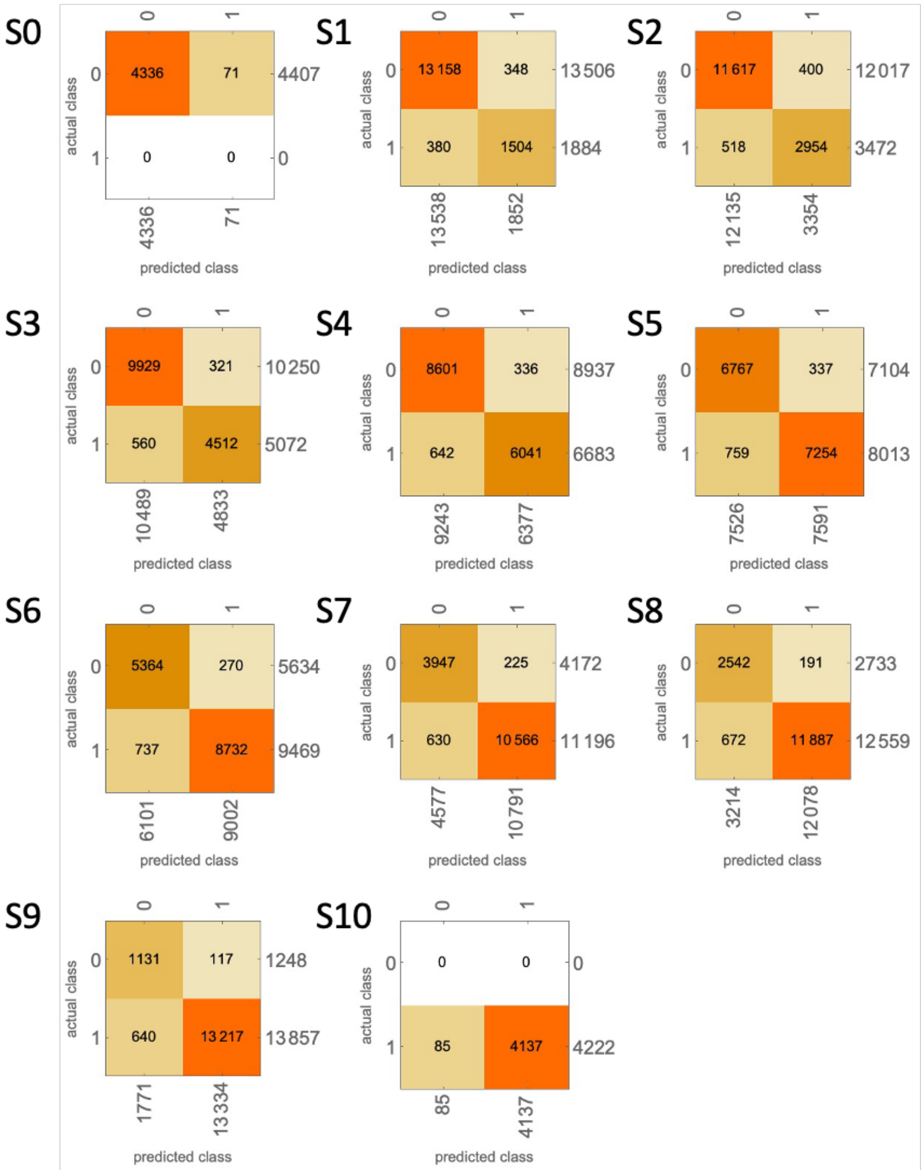


**Figure S5 Confusion matrices.** Shown are the confusion matrices for each dataset from the GBT model. Diagonal elements (top-left and bottom-right) represent correct classifications whereas off-diagonal elements (top-right and bottom-left) represent misidentification.

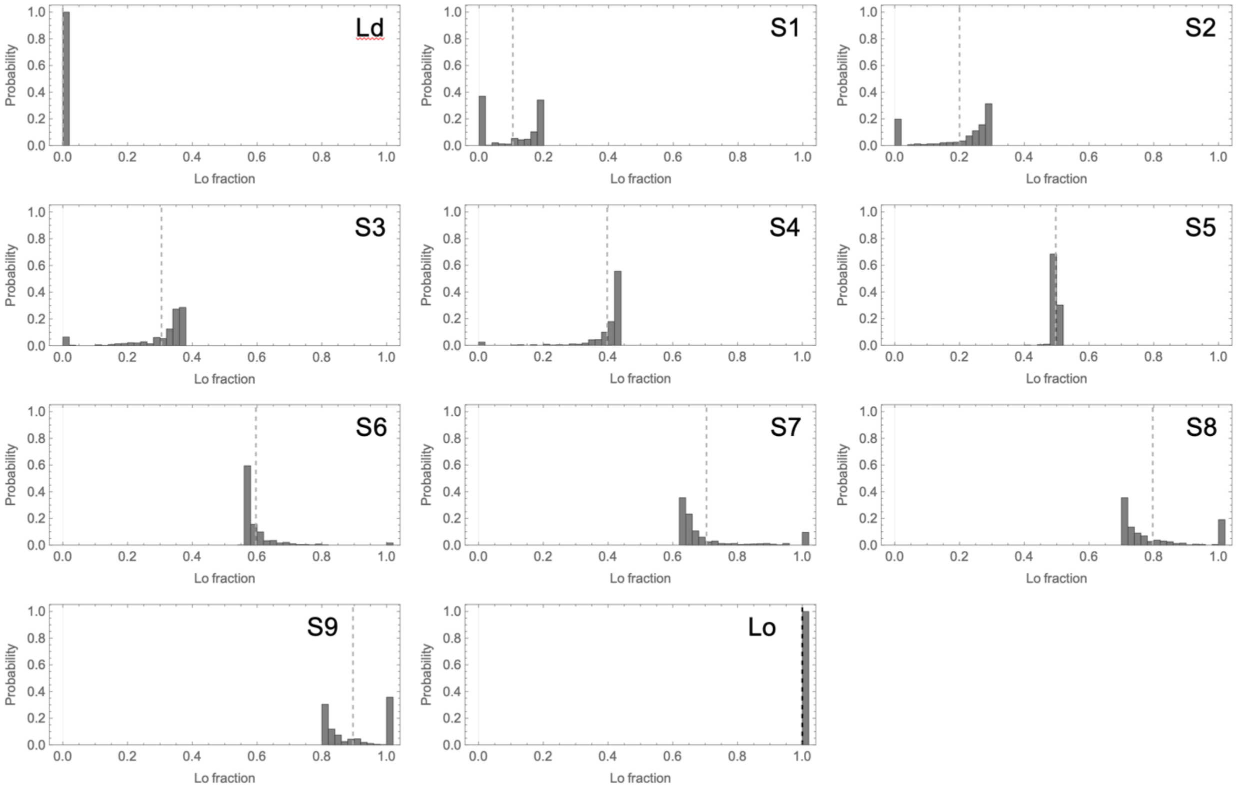


**Figure S6 Variability of apparent phase fraction for randomly oriented vesicles.** Shown are histograms of the ground truth Lo phase fraction for the 490 vesicles in each dataset as indicated in the figure. The mean Lo fraction is shown as a dashed line.

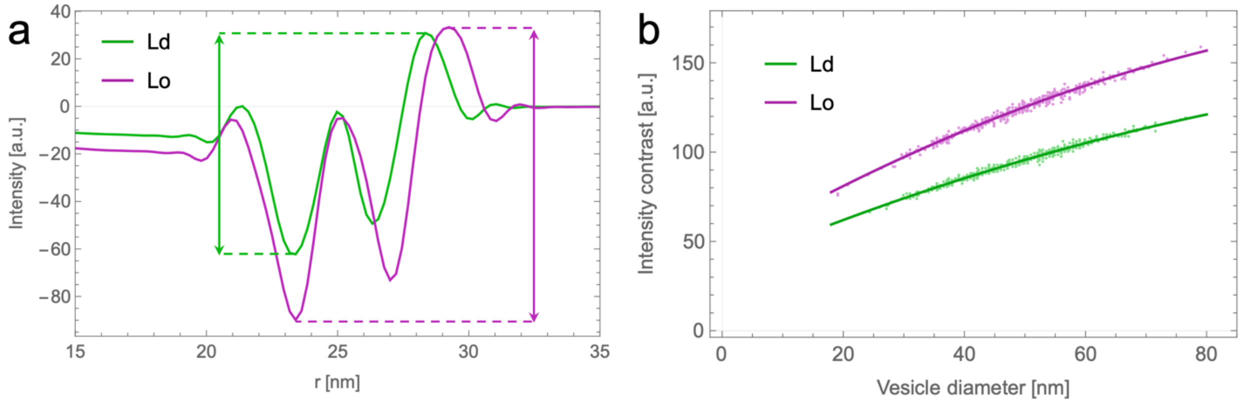


**Figure S7 Influence of vesicle size on intensity contrast.** (a) Ground truth IPs for Ld (green) and Lo (magenta) phases. Internal contrast is defined as the intensity difference between the outer peak and inner trough as indicated in the figure. (b) Scatter plot of internal contrast for the 490 vesicles of the Ld (green) and Lo (magenta) datasets. The solid lines are the best fit to a quadratic function.

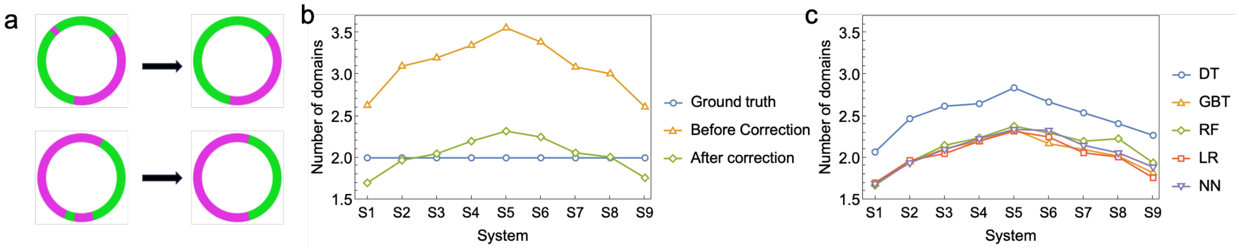


**Figure S8 Predicting the number of domains per vesicle.** (a) Schematic illustration of the correction used to eliminate isolated segments that are likely to be misclassified. (b) Number of domains calculated using LR before and after the correction, shown for each dataset. (c) Number of domains calculated using each supervised method, shown for each dataset.
